# The Genetic Legacy of Introgression from Two Deeply Divergent Lineages in the Genus *Homo*

**DOI:** 10.64898/2026.03.22.713509

**Authors:** Alan R. Rogers, Md Touhidul Islam, Colin M. Brand, Timothy H. Webster

## Abstract

Ancient DNA has shown that a distantly-related “superarchaic” population interbred first with the ancestors of Neanderthals and Denisovans and later with Denisovans themselves. Other work has shown that a superarchaic population interbred with the African ancestors of all modern humans. But it is not yet clear whether these events involved the same superarchaic population. Here, we use the distribution of derived alleles among populations to evaluate hypotheses about superarchaics and their relationship to other hominins of the Pleistocene and Holocene. We find evidence for at least two distinct superarchaic populations. The one contributing to archaic Eurasian populations (Denisovans and Neanderthal-Denisovan ancestors) diverged earlier from the human lineage than did the one contributing to early moderns in Africa. This latter superarchaic contributed a large fraction—about 20%—to the ancestry of modern humans. These findings reveal previously unrecognized structure among hominin populations in the Pleistocene.

**Significance statement:** Genetic data have shown that all human populations are mixtures. Since the last ice age, the populations mixing have been relatively close relatives—separated by a few tens of thousands of years or less. Earlier in the Pleistocene, there was mixing between more distantly-related populations, with separation times in excess of a million years. Here, we document admixture events involving two distinct populations, each of which had separated from the human lineage well over a million years ago. One of these was apparently Eurasian, because it mixed with archaic Eurasian populations. The other was apparently African, because it mixed with the African ancestors of modern humans. This latter event was remarkable, because the distant relative contributed such a large fraction—about 20%—to the ancestry of modern humans. These results show that during the Pleistocene, the earth was home to hominin populations far more distantly related than any pair of modern human populations, with separation times similar to that of bonobos and chimpanzees. These distantly-related pairs existed not only on separate continents, but also within the African continent. And in spite of the large differences between them, these populations were able to interbreed.

## Introduction

The past 15 years have revolutionized our understanding of interbreeding between ancient populations. There was interbreeding not only between different groups of modern humans, but also between moderns and archaic populations of Neanderthals and Denisovans. Interbreeding was documented not only in Eurasia, but also in African populations, where the source population was often obscure [9, 17, 21]. There was also interbreeding between Eurasian archaic populations and a distantly-related population, which diverged early in the Pleistocene [33, 34]. Because this population diverged earlier than did archaics, it has been called “superarchaic” [24, p. S167]. There is evidence for superarchaic gene flow into Denisovans [14, 24, 25], into Neanderthal-Denisovan ancestors [31], and into Africans [5]. More recently, several publications have argued that early modern humans were a mixed population that received a substantial fraction of ancestry from a superarchaic population that separated from the modern-archaic trunk over a million years ago [4, 8, 26, 35].

It is not yet clear whether there was one superarchaic population or several. Did the same superarchaic population contribute genes not only to archaic Eurasian populations but also to African ones? Or were there several such populations? In what follows, we present evidence for the second of these alternatives. We show that there was a second supearchaic population, which diverged later than the first and contributed ancestry not to archaic Eurasians, but to the African ancestors of modern humans. This implies that Pleistocene hominin populations had geographic structure much deeper than that of modern humans. There were deep separations, which stretched back over a million years, not only between populations on different continents, but also within Eurasia and within Africa.

## Results

### Comparing models of population history using nucleotide site patterns

Our notation is illustrated in Fig. 1, using a model proposed previously [31]. Capital letters refer to populations: *X* for west Africa, *Y* for Europe, *N* for Neanderthal, *D* for Denisovan, and *S* for superarchaic. Combinations of these letters refer to ancestral populations. For example, *XY* is the population ancestral to *X* and *Y*. Greek letters refer to episodes of admixture. The green, blue, and dotted lines indicate the gene genealogy of a hypothetical genetic locus at which we have observed one haploid sample each from populations *X, Y, N*, and *D*. Greek letters refer to episodes of gene flow. For example, *α* refers to *N → Y* gene flow—from population *N* into *Y*, and *δ* refers to *S → ND* gene flow. Combinations of Greek letters refer to models. For example, Fig. 1 illustrates model *αβγδ*, which includes four episodes of gene flow: *α, β, γ*, and *δ*.

**Figure 1:**
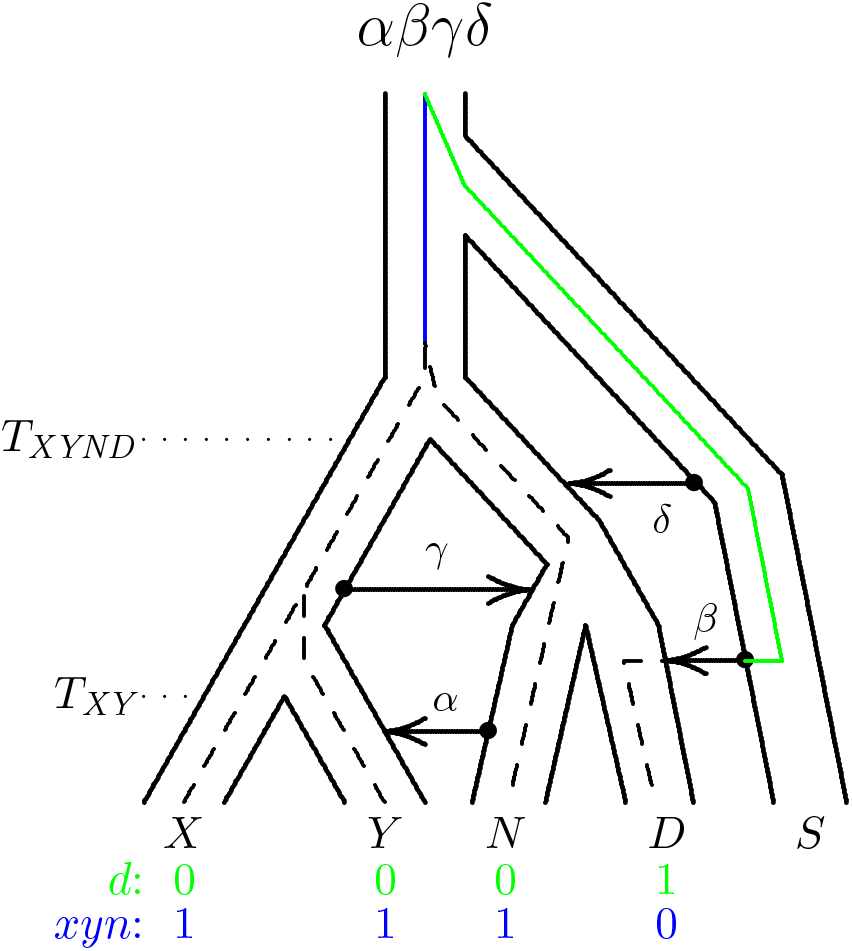
Model of population history proposed in 2020 [31]. Greek letters are episodes of gene flow; roman letters are populations: *X*, Africa; *Y*, Europe; *N*, Neanderthal; *D*, Denisovan; *S*, Superarchaic. *T*_*XY*_ is the separation time of *X* and *Y*, and *T*_*XYND*_ is that of *XY* (the population ancestral to *X* and *Y*) from *ND* (that ancestral to *N* and *D*). Internal lines show the gene genealogy of a single nucleotide site. A mutation on the green edge would generate site pattern *d*; one on blue would generate *xyn*.

Our analysis involves the frequencies of *site patterns*, two of which are shown on the bottom of the graph. If a mutation occurred on the green branch, the sample from *D* would carry the derived allele (“1”) at this locus, and the other samples would carry the ancestral allale (“0”). This configuration is called the “*d* site pattern” and is shown in green at the bottom of the graph. Similarly, a mutation on the blue branch would generate site pattern *xyn*, which is shown at the bottom in blue. The relative frequencies of site patterns can be estimated from data, and their expected values calculated using our *Legofit* software [29, 30]. Legofit estimates parameters by minimizing the difference between observed and expected site pattern frequencies.

We used 12 published high-coverage genomes, which we grouped into 4 sets: *X*, 3 genomes from the modern Yoruba of west Africa [19]; *Y*, 5 from the modern French and English [19]; *R*, 2 Neanderthals (Chagyrskaya [18] and Vindija [25]); *A*, 1 older Neanderthal (Altai [24]); and *D*, Denisovan [27]. Site patterns are labeled with combinations of these letters in lower case.

Fig. 2 shows the two most complex models considered in the current analysis. On the left, model *αβγδϵ* adds one episode of gene flow to the previous model (Fig. 1). In this episode, *ϵ*, early modern humans (*XY*) receive gene flow from the same superarchaic population that contributed to Denisovans and to the ancestors of Neanderthals and Denisovans. The model on the right (*αβγδζ*) also assumes that *XY* receives gene flow, but in this case the source is a distinct superarchaic population, *Z*, which diverged from the trunk more recently than *S*. In addition to these complex models, we also considered several simpler ones, which omit one or more episodes of gene flow.

**Figure 2:**
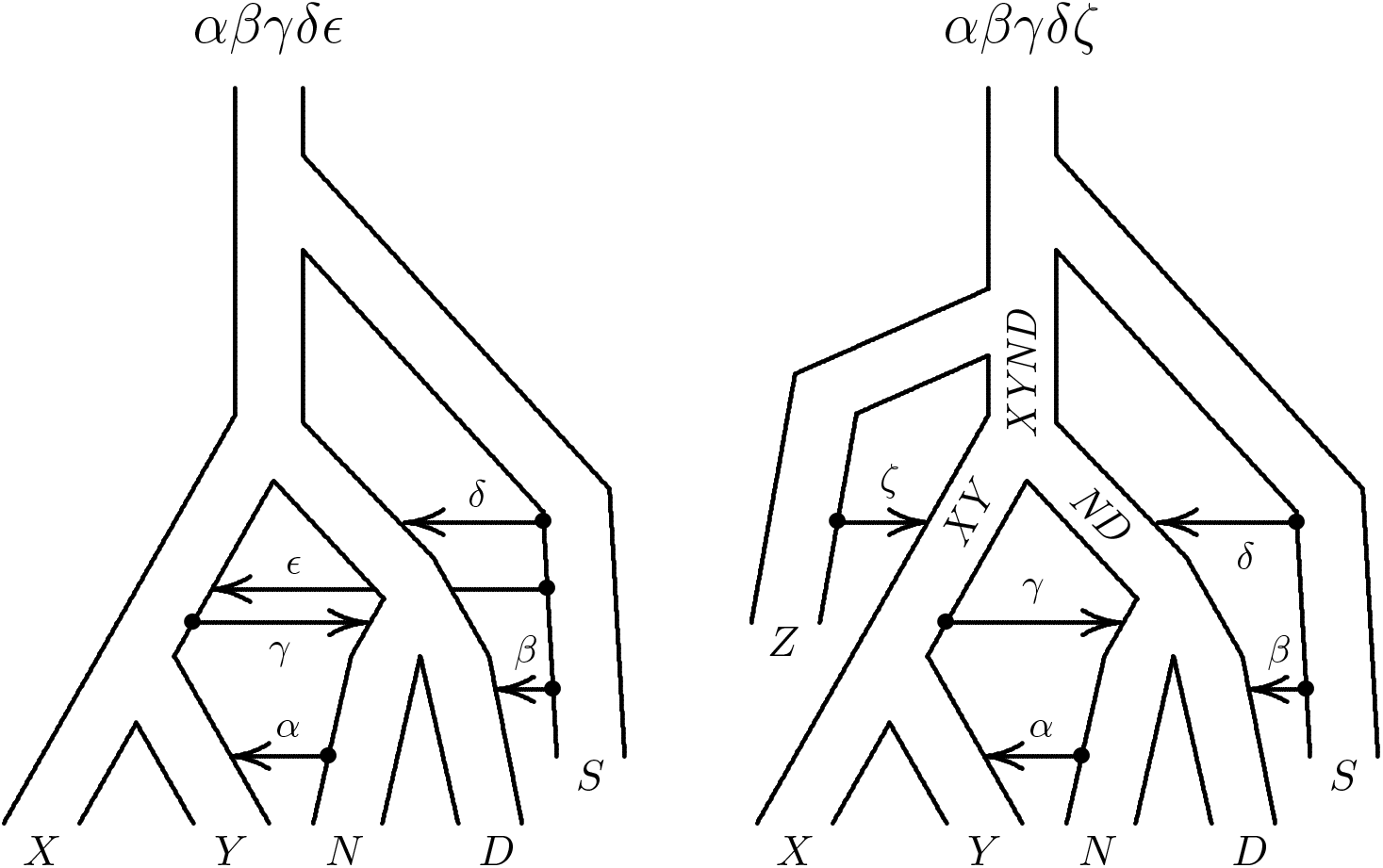
Two new models of population history with superarchaic admixture into the ancestors of modern humans. On the right, *Z* is a superarchaic population that diverged more recently than *S* and contributed ancestry (*ζ*) to ancestral moderns. *XY, ND*, and *XYND* label ancestral populations.

### Evidence for two distinct superarchaic populations

Our statistical analysis supports model *αβγδζ*, in which there are two distinct populations of superarchaics, one contributing to archaics and another contributing to the ancestors of modern humans. This conclusion rests on the “bootstrap estimate of predictive error” (bepe), a statistical method for choosing among models [6, 7]. Bepe uses the real data to estimate parameters and calculate expected site-pattern frequencies. It then compares these expectations to site-pattern frequencies in all 50 replicates produced by a moving-blocks bootstrap [16]. Bepe is the mean squared difference between predicted frequencies and those observed in bootstrap replicates (plus a correction for bootstrap bias). The preferred model is the one whose bepe value is lowest. In table 1, the preferred model is *αβγδζ*, but the other bepe values are also small. The question thus arises, how strongly should we weight the preferred model?

**Table 1:** Bootstrap estimate of predictive error (bepe) values and bootstrap model average (booma) weights.

| Model | bepe | weight |
| --- | --- | --- |
| $\alpha\beta\gamma$ | $2.17 \times 10^{-7}$ | 0 |
| $\alpha\beta\delta$ | $1.62 \times 10^{-7}$ | 0 |
| $\alpha\gamma\delta$ | $4.94 \times 10^{-7}$ | 0 |
| $\beta\gamma\delta$ | $6.06 \times 10^{-7}$ | 0 |
| $\alpha\beta\gamma\delta$ | $2.85 \times 10^{-7}$ | 0 |
| $\alpha\beta\gamma\zeta$ | $1.71 \times 10^{-7}$ | 0.02 |
| $\alpha\beta\delta\zeta$ | $1.84 \times 10^{-7}$ | 0 |
| $\alpha\gamma\delta\zeta$ | $2.62 \times 10^{-7}$ | 0 |
| $\alpha\beta\gamma\delta\epsilon$ | $1.46 \times 10^{-7}$ | 0 |
| $\alpha\beta\gamma\delta\zeta$ | $1.35 \times 10^{-7}$ | 0.98 |

The support for model *αβγδζ* is strong. This conclusion rests on the method of “bootstrap model averaging” [2] (booma). Booma iterates across all 51 data sets (the real data and 50 bootstrap replicates), pretends temporarily that the current data set is real and the others are bootstrap replicates, and calculates bepe for all models. The booma weight of each model is the fraction of these contests that it wins, i.e. the fraction in which its bepe value is lowest. Table 1 shows that model *αβγδζ* gets the preponderance of the booma weight. One other model gets 2% of the weight, and the rest get 0. Bootstrap replicates approximate repeated samples from the process that created the data. Consequently, these results imply that the advantage of the preferred model is large compared with variation in repeated sampling.

Of the two models in Fig. 2, the one on the left gets 0 weight, whereas the one on the right gets 98%. This is strong evidence that early modern humans received gene flow from a distinct superarchaic population—not from the one that contributed to Denisovans (episode *β*) and to the ancestors of Neanderthals and Denisovans (*δ*).

After fitting the preferred model, *αβγδζ*, the residual errors are generally small, as shown in Fig. 3. Nonetheless, several of them deviate significantly from 0, suggesting that the model is not yet a perfect fit. On the other hand, we don’t expect a perfect fit—any effort to select models by minimizing residuals would result in overfitting.

**Figure 3:**
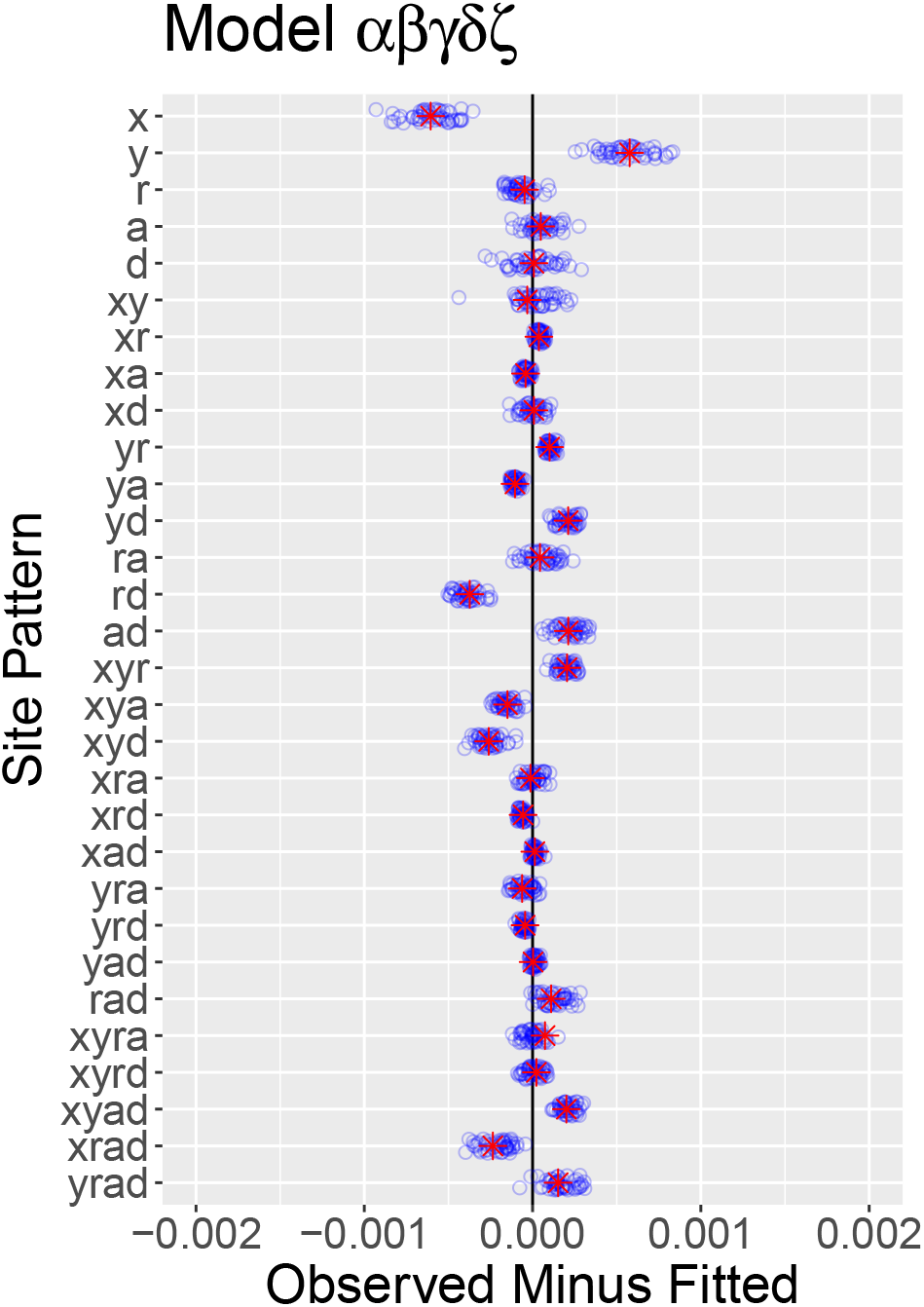
Residual errors in real data (red asterisks) and bootstrap replicates (blue circles)

Model-averaged parameter estimates (Fig. 4) are in general agreement with previous work [31], but there are a few exceptions. 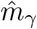 has changed from 0.016 to 0.045, implying more gene flow from early moderns into Neanderthals. 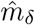 has changed from 0.034 to 0.058, implying more gene flow from Asian superarchaics into the Neanderthal-Denisovan ancestors. 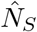 has changed from 27 k to 2.4 M, implying a larger size or deeper subdivision within the population of Asian superarchaics.

**Figure 4:**
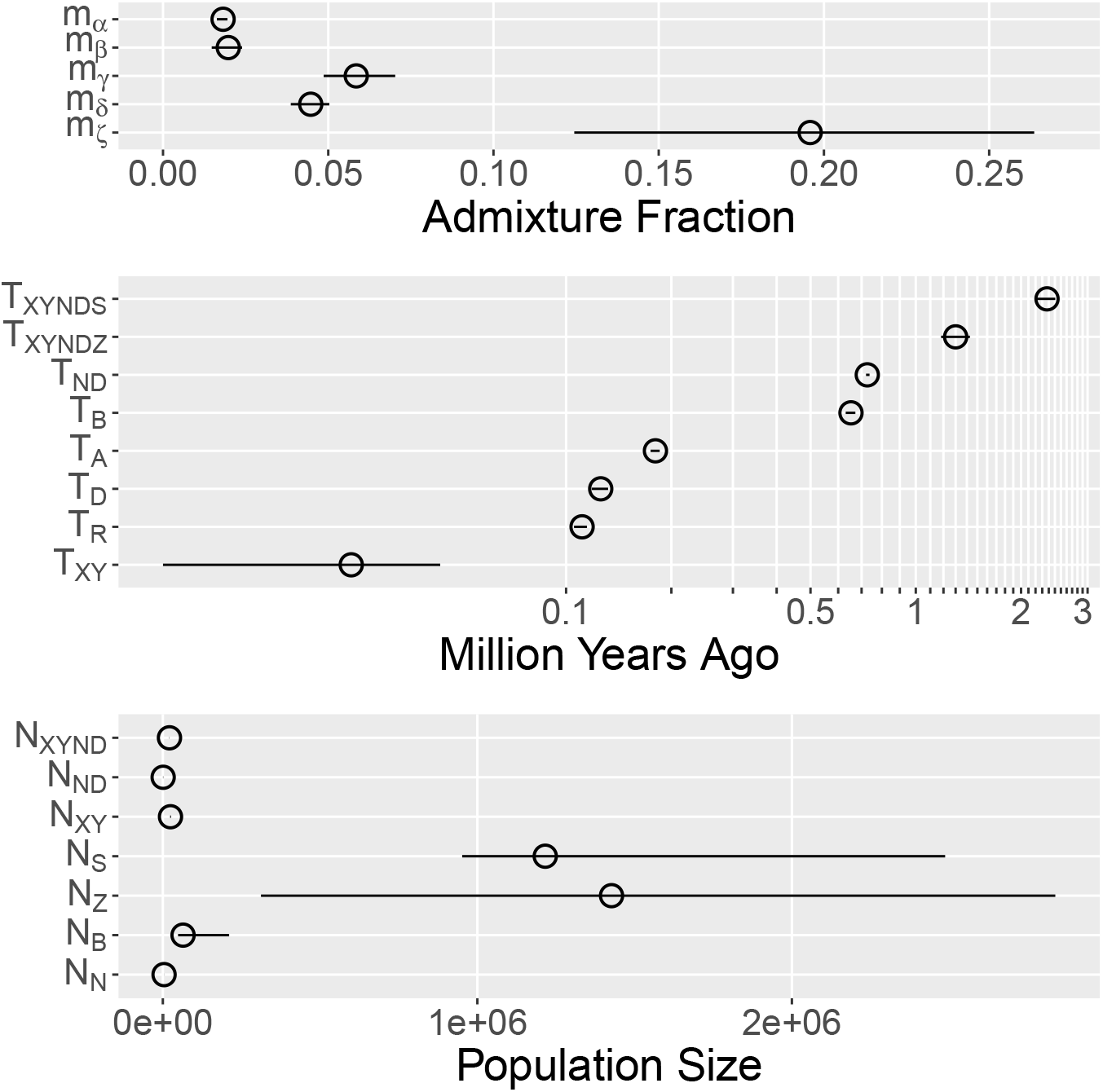
Model-averaged parameter estimates and confidence intervals estimated by moving-blocks bootstrap [16]. Admixture fractions: *m*_*i*_, fraction of recipient population derived from donor during admixture episode *i*. Time parameters: *T*_*XYNDS*_, separation time of *S* from the ancestors of *X, Y, N*, and *D*; *T*_*XYNDZ*_, separation time of *Z* from the ancestors of *X, Y, N*, and *D*; *T*_*ND*_, separation time of *N* and *D*; *T*_*B*_, time at which the Neanderthal population changed size; *T*_*A*_, age of the Altai Neanderthal; *T*_*D*_, age of the Denisovan genome; *T*_*R*_, average age of the Chagyrskaya and Vindija Neanderthals; *T*_*XY*_, separation time of *X* and *Y*. Effective population sizes: *N*_*XYND*_, size of population ancestral to *X, Y, N*, and *D*; *N*_*ND*_; size of population ancestral to *N* and *D*; *N*_*XY*_, size of population ancestral to *X* and *Y*; *N*_*S*_, size of population *S*; *N*_*Z*_, size of population *Z*; *N*_*B*_, size of early Neanderthal population; *N*_*N*_, size of later Neanderthal population.

Although many of the confidence intervals in Fig. 4 are narrow, some are quite large, and several of these large intervals reflect identifiability problems. For example, supplementary Fig. S3 shows that estimates of *N*_*Z*_ and *N*_*B*_ cluster tightly along a straight line with positive slope. This implies that composite likelihood depends on these parameters primarily through their difference, so that an increase in one parameter can be offset by an increase in the other.

Identifiability problems such as these can underlie not only wide confidence intervals but also bias. Our legofit analysis may have additional bias, because it ignores states with probability less than 10^*−*6^, as discussed below. To measure these biases, we analyzed 50 data sets simulated with msprime [12] under the preferred model, *αβγδζ*. Fig. 5 shows the results. In this figure, red crosses are true parameters values used as input by the simulation, and blue circles are legofit estimates. Admixture and time parameters show little bias, although there is substantial variation among estimates of *m*_*ζ*_ and *T*_*XY*_. Among population size parameters with large values (*N*_*B*_, *N*_*Z*_, and *N*_*S*_) most estimates are larger than the true values, indicating an upward bias. These parameters describe the sizes of the early Neanderthal population and of the two superarchaic populations. Consequently, the estimates in Fig. 4 may be too large. Among size parameters with smaller values (*N*_*N*_, *N*_*XY*_, *N*_*ND*_, and *N*_*XYND*_), the opposite is true—estimates are biased downward, and the real estimates in Fig. 4 are probably too small. In spite of these problems, these results show that in most cases, the true values are either close to their estimates or are surrounded by a relatively unbiased swarm. Thus, legofit does a reasonable job of fitting this model.

**Figure 5:**
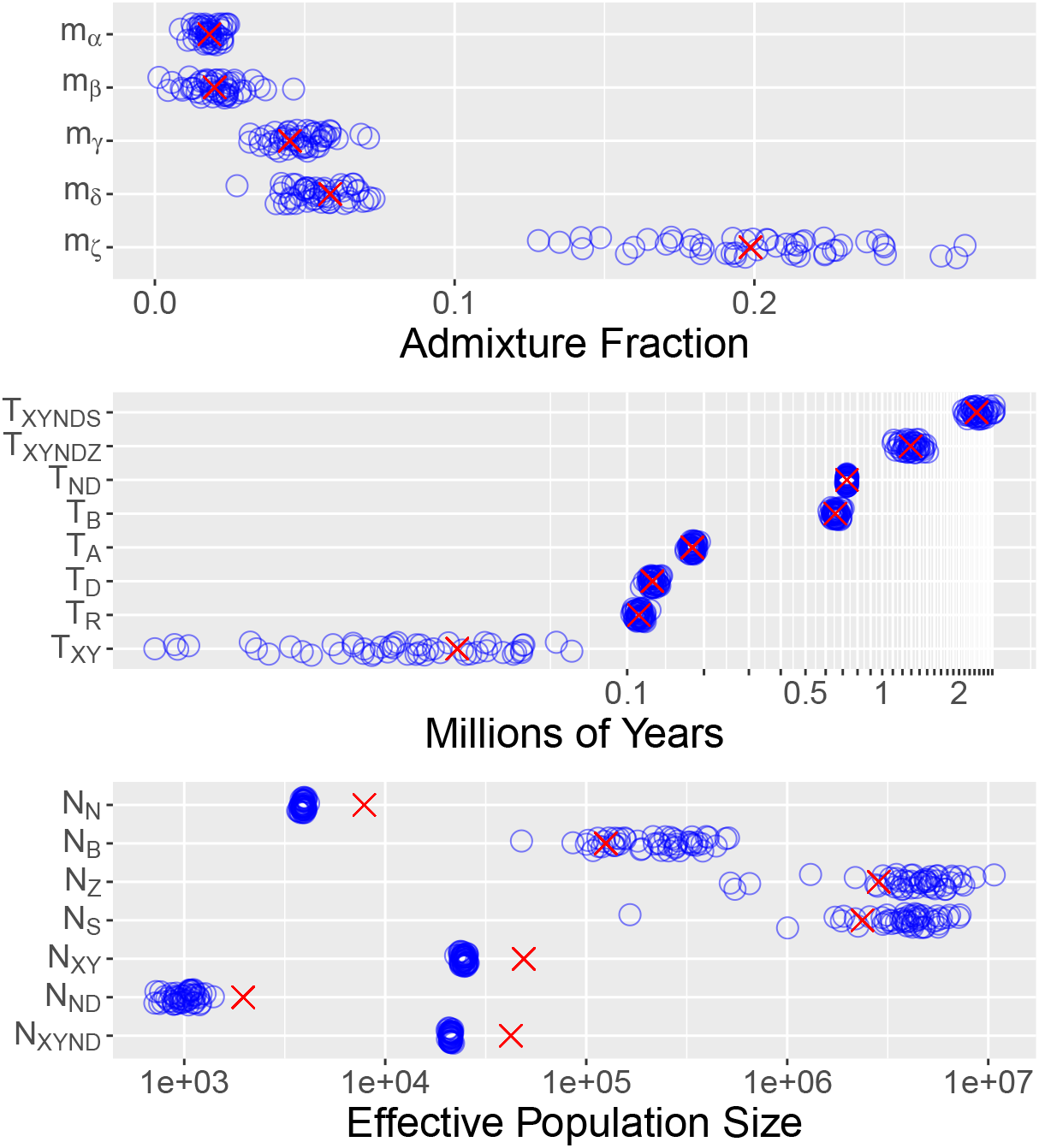
Estimates from simulations of the fitted model. Blue circles show estimates based 50 data sets simulated with msprime [12]. Red crosses show true parameter values. Parameters are defined in the caption of Fig. 4.

## Discussion

These results solve a puzzle: under model *αβγδ* [31] (Fig. 1), successive methodological improvements led to less and less plausible estimates of the separation time, *T*_*XY*_, between modern west Africans and Europeans. This separation probably happened about 50 ka ago, a date supported both by archaelogical and genetic data [e.g. 13, 32]. Our first estimate of this parameter (row 1, supplementary table S2) is consistent with this consensus. The table’s second row differs from the first only in using a more accurate algorithm [30]. The third row adds an additional archaic genome [18]. With each improvement, the estimate of *T*_*XY*_ becomes less plausible. By row 3 it has become absurd, implying that Europeans and Africans separated only a few thousand years ago. This suggests that the model is misspecified, resulting in a bias that becomes clearer with greater accuracy and more data.

This bias arises because Africans and Europeans uniquely share more derived alleles than model *αβγδ* can easily accomodate. In response, the optimization algorithm wants to increase the length of the branch ancestral to west Africans and Eurasians in the population tree. In principal, it could do so either by (1) increasing the separation time, *T*_*XYND*_, of moderns and archaics, or by (2) reducing the separation time, *T*_*XY*_, of Eurasians and Africans. (These parameters are illustrated in Fig. 1.) Option 1 is unavailable however because *T*_*XYND*_ has a fixed value that is used to calibrate the molecular clock. Thus, only *T*_*XY*_ is free to vary, and the optimizer has pushed it to an implausibly low value. The present results resolve this discrepancy. In the model-averaged estimates, the upper end of the *T*_*XY*_ confidence interval is about 44 ka, which is not far short of accepted views about the separation time of Europeans and west Africans.

Our results agree with several recent publications [4, 8, 36], which find that the early modern human population (*XY* in the right panel of Fig. 2) received substantial gene flow from a superarchaic population (our *Z*) that diverged prior to the separation of archaics from the ancestors of moderns. Population *Z* separated from *XYND* 1–3 Ma ago in the reconstruction of Fan et al. [8, p. 929], ~1.5 Ma in that of Cousins et al. [4, p. 858], over 500 ka in that of Zhang et al. [36] and Ma (95% CI: 1.181–1.428) in ours, assuming generations of 29 y. The contribution of population *Z* is 5–15% for Fan et al., 21% for Cousins et al., 0.5–1.1% for Zhang et al., and 19.6% (95% CI: 12.4–26.4%) for us.

In another recent study, Ragsdale et al. [26] agree that early modern humans were a mixture and that the population contributing the majority to this mixture is related to Neanderthals and Denisovans. But they find that the fit is improved if continuous gene flow is allowed between the two populations contributing to this mixture. Perhaps because of this gene flow, their estimate of the separation time is larger—about 1.7 Ma ago.

Some African samples carry a long haplotype at the MUC7 locus [35]. The large difference between this haplotype and others indicates that it separated several million years ago. Yet its length indicates that it arrived in the modern human population only recently. This evidence also supports the view that modern Africans carry DNA derived by interbreeding with a superarchaic population.

The similarity between these sets of findings is remarkable in view of the differences between methods. Some of the methods ignore linkage disequilibrium whereas others rely on it. Some use variation within populations (or even within individuals) whereas others use differences between populations. Yet the studies reach similar conclusions and are thus mutually supportive.

Our own study is unique in showing that archaics and moderns interbred with two distinct superarchaic populations—one contributing to archaics and another to early moderns. This new finding requires revision of our previous views about the Middle Pleistocene. In an earlier publication [31], we found that the ancestors of Neanderthals and Denisovans interbred with the same superarchaic population that later interbred with Denisovans. To make sense of this, we pointed out that the first wave of emigration out of Africa happened early in the Pleistocene, when *Homo erectus* spread across Eurasia. Later, during the Middle Pleistocene, humans evolved larger brains and began making Acheulean tools. Both of these innovations appear in Africa before Eurasia, suggesting a second wave of emigration out of Africa [10, 11]. We proposed [31] that this second wave interbred with the Eurasian descendants of the first, during what we refer to as the *δ* episode of admixture. It seemed plausible that, before this contact, the two populations had remained largely isolated because of the difficulty of traveling between Africa and Eurasia. (At least during the Upper Pleistocene, human contact between these continents was largely restricted to relatively brief periods when the Sahara was humid [3].)

Now we have evidence of a second superarchaic population, *Z*, which diverged after the first and later interbred with the ancestors of modern humans. This contact presumably occurred in Africa, because it happened before moderns spread into Eurasia. We considered the possibility that this second superarchaic population was the same as the first—see model *αβγδϵ* in Fig. 2—but that model fared poorly (table 1). Thus, there were two superarchaic populations, and it seems likely that the second (population *Z*) was African.

This is puzzling, because it implies that two African populations—population *Z* and the ancestors of moderns, Neanderthals, and Denisovans—remained essentially isolated across roughly a million years. What kept them apart? Africa has no mountain barriers as large as the Himalayas or the Alps. There are deserts, but these were not continuously arid [20]. The results of Ragsdale et al. [26] (see above) suggest that these populations may not have been isolated after all—perhaps there was a continuous trickle of gene flow between them. If this gene flow did occur, it would cause a downward bias in other estimates of the separation time of populations *Z* and *XYND* —not only our estimates, but also those of Fan et a. [8, p. 929] and Cousins et al. [4, p. 858]. Yet these estimates are all well over a million years, and this suggests that any gene flow between *Z* and *XYND* must have been weak.

In broader perspective, we can distinguish three epochs of admixture among Pleistocene hominins. During the past 20 ky, there is evidence of extensive mixing between populations of modern humans [28]. In this most recent epoch, the populations mixing were not distantly related, having separated no more than a few tens of thousands of years earlier. The next earlier epoch of mixing was around 40 ky ago, as modern humans expanded out of Africa and mixed with archaic populations of Neanderthals and Denisovans. During this epoch, there was mixing between populations that were more distantly related, having been separate for about half a million years. Prior to this, there was mixing between populations that had been separate for over a million years—similar to the separation of bonobos and chimpanzees [23, p. 529, 1].

These facts reveal an interesting change in global population structure. A few hundred thousand years ago, the world was home to hominin populations that were as different from each other as are modern bonobos and chimpanzees. These large differences occurred not only between populations on different continents, but also within Africa. And in spite of these differences, it was possible for at least some of these distant relatives to interbreed.

By forty thousand years ago, population differences had shrunk but were still larger—with separations of a half-million years—than any seen today. During the last 20 ky, population differences have been similar to modern ones.

The homogeneity of modern human populations is easy to understand as a consequence of mixing and population replacements. But why were earlier differences so large? Perhaps geographic barriers, such as deserts, mountain ranges, and rivers, were more salient then, just as they are to modern bonobos and chimpanzees. Or maybe rates of local extinction and recolonization were lower. The causes of any such differences are as yet unknown.

## Methods

We used 12 published high-coverage genomes grouped into 5 subsets: subset *X* consists of the 3 Yoruban genomes published by the Simons Genome Diversity Project [19] (SGDP); subset *Y* consists all 3 French and both English samples in the SGDP; subset *R* consists of 2 relatively recent Neanderthal genomes (Vindija [25] and Chagyrskaya [18]); subset *A* is the Altai Neanderthal genome [24]); and subset *D* is the high-coverage Denisovan genome [27].

All genomes were downloaded from the websites of the SGDP and the Max Planck Institute for Evolutionary Anthropology. Methods for quality control, calling ancestral alleles, and calibrating the molecular clock are as described by Rogers et al. [31].

Legofit [29, 30] estimates parameters by minimizing the Kullback-Leibler (KL) divergence [15] between observed and expected site-pattern frequencies. Its minimization engine uses the differential evolution [22] (DE) algorithm, which maintains a swarm of points—10 times as many as there are parameters—each of which represents a set of guesses about all free parameters. In each iteration, the points with lowest KL divergence mutate, recombine, and reproduce to form the next generation of points.

Our Legofit analysis involves four stages. The first two are very fast and achieve only an approximate fit. Then principal components (PC) analysis reduces the number of parameters and re-expresses free variables on orthogonal coordinates. Finally, legofit’s deterministic algorithm provides a more accurate fit. Analysis of most models was as follows:

1. Stage a1 used legofit’s stochastic algorithm with arguments --tol 3e-5 -S 100@1000 -S 1000@10000. This says that the tolerance criterion is 3e-5, which is fairly large and means that the DE iterations will stop when the fit is only fair. The other arguments say that we start with 100 DE iterations in each of which the objective function is estimated from 1000 simulation replicates. Then it does 1000 DE iterations with 10000 replicates per function evaluation.
2. Stage a2 also uses stochastic algorithm and uses the same `--tol` value as a1. It initializes the DE algorithm with points sampled from the point swarms of all data sets (real data and bootstrap replicates) in stage a1. In this stage, legofit does at most 500 DE iterations with 10000 replicates per function evaluation.
3. Stages b1 and b2 operate on principal components (PCs) rather than on the original variables. PCs that account for less than a fraction 0.001 of the total variance are ignored. Stage b2 also does 500 DE iterations of the stochastic algorithm with 10000 replicates per function evaluation.
4. Stage b2 initializes the DE algorithm by sampling from the point swarms of all data sets in stage b1. It uses the deterministic algorithm with argument `-d 1e-6`, which says to ignore states with probability less then 1e-6. Without this approximation, the deterministic algorithm would be not be feasible with our most complex models. The summed probability of ignored events was a fraction 0.0008 of the total probability. The tolerance criterion is dropped to `--tol 3e-6` for this final stage and there are 2000 DE iterations. In no case did the algorithm go the full 2000 iterations. It always stopped early because it had reached the tolerance criterion.

We don’t estimate the timing of admixture events, because most of the information about these parameters is in linkage disequilibrium, which our method ignores. Because our method is insensitive to these parameters, we constrain their values to arbitrary time points within the relevant interval. For example, *T*_*β*_ is constrained to equal (*T*_*D*_ + *T*_*ND*_)*/*2. Some episodes of admixture (*α* and *β*) affect only one sampled genome. For these episodes, the timing of admixture has no effect on likelihood, and we lose nothing by fixing the values of these parameters. The other episodes (*γ, δ*, and *ζ*) affect more than one sampled genome, and for these our approach is an approximation.

In some of the smaller models (*αβγ, αβδ, αγδ, βγδ*), which were done early in the project, we used the derministic algorithm with `-d 0.` This does not ignore events of low probability and thus makes the calculations more accurate. Ordinarily we wouldn’t compare models fit using different values of the `-d` parameter, because smaller values provide greater accuracy and give the model an advantage in model selection. But since these small models were all excluded in model selection, we did not re-run these models using the same parameters used for the large ones.

## Acknowledgements

This work was supported by NSF BCS 1945782 (ARR and TWH) and the Center for High Performance Computing at the University of Utah (ARR). All scripts and pipelines are available in file `Ancestry-from-2-superarchaic-populations.tgz` at https://doi.org/10.6084/m9.figshare.31880014.

## Author contributions

ARR developed the Legofit software, did Legofit analyses, and wrote the manuscript. MTI did Legofit analyses. All authors collaborated on study design, interpretation, and manuscript revision.

## References

[1] Colin M. Brand, Frances J. White, Alan R. Rogers, and Timothy H. Webster (2022). “Estimating bonobo (Pan paniscus) and chimpanzee (Pan troglodytes) evolutionary history from nucleotide site patterns”. Proceedings of the National Academy of Sciences, USA 119.17, e2200858119. doi: 10.1073/pnas.2200858119.

[2] Steven T Buckland, Kenneth P Burnham, and Nicole H Augustin (1997). “Model selection: an integral part of inference”. Biometrics 53.2, pp. 603–618. doi: 10.2307/2533961.

[3] Isla S. Castañeda, Stefan Mulitza, Enno Schefuß, Raquel A. Lopes dos Santos, Jaap S. Sinninghe Damsté, and Stefan Schouten (2009). “Wet phases in the Sahara/Sahel region and human migration patterns in North Africa”. Proceedings of the National Academy of Sciences, USA 106.48, pp. 20159–20163. doi: 10.1073/pnas.0905771106.

[4] Trevor Cousins, Aylwyn Scally, and Richard Durbin (Apr. 2025). “A structured coalescent model reveals deep ancestral structure shared by all modern humans”. Nature Genetics 57.4, pp. 856–864. doi: 10.1038/s41588-025-02117-1.

[5] Arun Durvasula and Sriram Sankararaman (2020). “Recovering signals of ghost archaic introgression in African populations”. Science Advances 6.7, aax5097. doi: 10.1126/sciadv.aax50.

[6] Bradley Efron (1983). “Estimating the error rate of a prediction rule: improvement on cross-validation”. Journal of the American Statistical Association 78.382, pp. 316–331. doi: 10.1080/01621459.1983.10477973.

[7] Bradley Efron and Robert J. Tibshirani (1993). An Introduction to the Bootstrap. New York: Chapman and Hall, 1993. doi: 10.1007/978-1-4899-4541-9.

[8] Shaohua Fan, Jeffrey P Spence, Yuanqing Feng, Matthew EB Hansen, Jonathan Terhorst, Marcia H Beltrame, Alessia Ranciaro, Jibril Hirbo, William Beggs, Neil Thomas, Thomas Nyambo, Sununguko Wata Mpoloka, Gaonyadiwe George Mokone, Alfred K. Njamnshi, Charles Fokunang, Dawit Wolde Meskel, Gurja Belay, Yun S. Song, and Sarah A. Tishkoff (2023).“Whole-genome sequencing reveals a complex African population demographic history and signatures of local adaptation”. Cell 186.5, pp. 923–939. doi: 10.1016/j.cell.2023.01.042.

[9] PingHsun Hsieh, August E Woerner, Jeffrey D Wall, Joseph Lachance, Sarah A Tishkoff, Ryan N Gutenkunst, and Michael F Hammer (2016). “Model-based analyses of whole-genome data reveal a complex evolutionary history involving archaic introgression in Central African Pygmies”. Genome Research 26.3, pp. 291–300.

[10] Jean-Jacques Hublin (1998). “Climatic changes, paleogeography, and the evolution of the Neandertals”. In: Neandertals and Modern Humans in Western Asia. Ed. by Takeru Akazawa, Kenichi Aoki, and Ofer Bar-Yosef. Kluwer, 1998, pp. 295–310.

[11] Jean-Jacques Hublin (2009). “The origin of Neandertals”. Proceedings of the National Academy of Sciences, USA 106.38, pp. 16022–16027.

[12] Jerome Kelleher, Alison M Etheridge, and Gilean McVean (May 2016). “Efficient coalescent simulation and genealogical analysis for large sample sizes”. PLoS Computational Biology 12.5, pp. 1–22. doi: 10.1371/journal.pcbi.1004842.

[13] Elise Kerdoncuff, Laurits Skov, Nick Patterson, Joyita Banerjee, Pranali Khobragade, Sankha S. Chakrabarti, Avinash Chakrawarty, Prasun Chatterjee, Minakshi Dhar, Monica Gupta, John P. John, Parvaiz A. Koul, Sarabmeet S. Lehl, Rashmi R. Mohanty, Mekala Padmaja, Arokiasamy Perianayagam, Chhaya Rajguru, Lalit Sankhe, Arunansu Talukdar, Mathew Varghese, Sathyanarayana Raju Yadati, Wei Zhao, Yuk Yee Leung, Gerard D. Schellenberg, Yi Zhe Wang, Jennifer A. Smith, Sharmistha Dey, Andrea Ganna, Aparajit Ballav Dey, Sharon L.R. Kardia, Jinkook Lee, and Priya Moorjani (June 2025). “50,000 years of evolutionary history of India: impact on health and disease variation”. Cell 188.13, 3389–3404.e6. doi: 10.1016/j.cell.2025.04.027.

[14] Martin Kuhlwilm, Ilan Gronau, Melissa J. Hubisz, Cesare de Filippo, Javier Prado-Martinez, Martin Kircher, Qiaomei Fu, Hernán A. Burbano, Carles Lalueza-Fox, Marco de la Rasilla, Antonio Rosas, Pavao Rudan, Dejana Brajkovic, Željko Kucan, Ivan Gušic, Tomas Marques-Bonet, Aida M. Andrés, Bence Viola, Svante Pääbo, Matthias Meyer, Adam Siepel, and Sergi Castellano (Feb. 2016). “Ancient gene flow from early modern humans into Eastern Neanderthals”. Nature 530.7591, pp. 429–433. issn: 1476-4687. doi: 10.1038/nature16544.

[15] Solomon Kullback and Richard A Leibler (Mar. 1951). “On information and sufficiency”. The Annals of Mathematical Statistics 22.1, pp. 79–86. doi: 10.1214/aoms/1177729694.

[16] Regina Y. Liu and Kesar Singh (1992). “Moving blocks jacknife and bootstrap capture weak dependence”. In: Exploring the “Limits” of the Bootstrap. Ed. by Raoul LePage and Lynne Billard. New York: Wiley, 1992, pp. 225–248.

[17] Belen Lorente-Galdos, Oscar Lao, Gerard Serra-Vidal, Gabriel Santpere, Lukas FK Kuderna, Lara R Arauna, Karima Fadhlaoui-Zid, Ville N Pimenoff, Himla Soodyall, Pierre Zalloua, Tomas Marques-Bonet, and David Comas (2019). “Whole-genome sequence analysis of a pan African set of samples reveals archaic gene flow from an extinct basal population of modern humans into sub-Saharan populations”. Genome Biology 20.1, p. 77.

[18] Fabrizio Mafessoni, Steffi Grote, Cesare de Filippo, Viviane Slon, Kseniya A. Kolobova, Bence Viola, Sergey V. Markin, Manjusha Chintalapati, Stephane Peyrégne, Laurits Skov, Pontus Skoglund, Andrey I. Krivoshapkin, Anatoly P. Derevianko, Matthias Meyer, Janet Kelso, Benjamin Peter, Kay Prüfer, and Svante Pääbo (2020). “A high-coverage Neandertal genome from Chagyrskaya Cave”. Proceedings of the National Academy of Sciences, USA 117.26, pp. 15132–15136. doi: 10.1073/pnas.2004944117.

[19] Swapan Mallick, Heng Li, Mark Lipson, Iain Mathieson, Melissa Gymrek, Fernando Racimo, Mengyao Zhao, Niru Chennagiri, Susanne Nordenfelt, Arti Tandon, Pontus Skoglund, Iosif Lazaridis, Sriram Sankararaman, Qiaomei Fu, Nadin Rohland, Gabriel Renaud, Yaniv Erlich, Thomas Willems, Carla Gallo, Jeffrey P. Spence, Yun S. Song, Giovanni Poletti, Francois Balloux, George van Driem, Peter de Knijff, Irene Gallego Romero, Aashish R. Jha, Doron M. Behar, Claudio M. Bravi, Cristian Capelli, Tor Hervig, Andres Moreno-Estrada, Olga L. Posukh, Elena Balanovska, Oleg Balanovsky, Sena Karachanak-Yankova, Hovhannes Sahakyan, Draga Toncheva, Levon Yepiskoposyan, Chris Tyler-Smith, Yali Xue, M. Syafiq Abdullah, Andres Ruiz-Linares, Cynthia M. Beall, Anna Di Rienzo, Choongwon Jeong, Elena B. Starikovskaya, Ene Metspalu, Jüri Parik, Richard Villems, Brenna M. Henn, Ugur Hodoglugil, Robert Mahley, Antti Sajantila, George Stamatoyannopoulos, Joseph T. S. Wee, Rita Khusainova, Elza Khusnutdinova, Sergey Litvinov, George Ayodo, David Comas, Michael F. Hammer, Toomas Kivisild, William Klitz, Cheryl A. Winkler, Damian Labuda, Michael Bamshad, Lynn B. Jorde, Sarah A. Tishkoff, W. Scott Watkins, Mait Metspalu, Stanislav Dryomov, Rem Sukernik, Lalji Singh, Kumarasamy Thangaraj, Svante Pääbo, Janet Kelso, Nick Patterson, and David Reich (2016). “The Simons Genome Diversity Project: 300 genomes from 142 diverse populations”. Nature 538, pp. 201–206. issn: 1476-4687. doi: 10.1038/nature18964.

[20] J von der Meden, R Pickering, BJ Schoville, H Green, R Weij, J Hellstrom, et al. (2022). “Tufas indicate prolonged periods of water availability linked to human occupation in the southern Kalahari”. PLoS ONE 17.7, e0270104. doi: 10.1371/journal.pone.0270104.

[21] Vincent Plagnol and Jeffrey D Wall (2006). “Possible ancestral structure in human populations”. PLoS Genetics 2.7, e105.

[22] Kenneth Price, Rainer M Storn, and Jouni A Lampinen (2006). Differential Evolution: A Practical Approach to Global Optimization. Berlin: Springer Science and Business Media, 2006. isbn: 978-3-540-20950-8.

[23] Kay Prüfer, Kasper Munch, Ines Hellmann, Keiko Akagi, Jason R. Miller, Brian Walenz, Sergey Koren, Granger Sutton, Chinnappa Kodira, Roger Winer, James R. Knight, James C. Mullikin, Stephen J. Meader, Chris P. Ponting, Gerton Lunter, Saneyuki Higashino, Asger Hobolth, Julien Dutheil, Emre Karakoç, Can Alkan, Saba Sajjadian, Claudia Rita Catacchio, Mario Ventura, Tomas Marques-Bonet, Evan E. Eichler, Claudine André, Rebeca Atencia, Lawrence Mugisha, Jörg Junhold, Nick Patterson, Michael Siebauer, Jeffrey M. Good, Anne Fischer, Susan E. Ptak, Michael Lachmann, David E. Symer, Thomas Mailund, Mikkel H. Schierup, Aida M. Andrés, Janet Kelso, and Svante Pääbo (June 2012). “The bonobo genome compared with the chimpanzee and human genomes”. Nature 486.7404, pp. 527–531. doi: 10.1038/nature11128.

[24] Kay Prüfer, Fernando Racimo, Nick Patterson, Flora Jay, Sriram Sankararaman, Susanna Sawyer, Anja Heinze, Gabriel Renaud, Peter H Sudmant, Cesare de Filippo, Heng Li, Swapan Mallick, Michael Dannemann, Qiaomei Fu, Martin Kircher, Martin Kuhlwilm, Michael Lachmann, Matthias Meyer, Matthias Ongyerth, Michael Siebauer, Christoph Theunert, Arti Tandon, Priya Moorjani, Joseph Pickrell, James C. Mullikin, Samuel H. Vohr, Richard E. Green, Ines Hellmann, Philip L. F. Johnson, Hélène Blanche, Howard Cann, Jacob O. Kitzman, Jay Shendure, Evan E. Eichler, Ed S. Lein, Trygve E. Bakken, Liubov V. Golovanova, Vladimir B. Doronichev, Michael V. Shunkov, Anatoli P. Derevianko, Bence Viola, Montgomery Slatkin, David Reich, Janet Kelso, and Svante Pääbo (2014). “The complete genome sequence of a Neanderthal from the Altai Mountains”. Nature 505.7481, pp. 43–49. doi: 10.1038/nature12886.

[25] Kay Prüfer, Cesare de Filippo, Steffi Grote, Fabrizio Mafessoni, Petra Korlević, Mateja Hajdinjak, Benjamin Vernot, Laurits Skov, Pinghsun Hsieh, Stéphane Peyrégne, David Reher, Charlotte Hopfe, Sarah Nagel, Tomislav Maricic, Qiaomei Fu, Christoph Theunert, Rebekah Rogers, Pontus Skoglund, Manjusha Chintalapati, Michael Dannemann, Bradley J. Nelson, Felix M. Key, Pavao Rudan, Željko Kućan, Ivan Gušić, Liubov V. Golovanova, Vladimir B. Doronichev, Nick Patterson, David Reich, Evan E. Eichler, Montgomery Slatkin, Mikkel H. Schierup, Aida Andrés, Janet Kelso, Matthias Meyer, and Svante Pääbo (2017). “A high-coverage Neandertal genome from Vindija Cave in Croatia”. Science 358.6363, pp. 655–658. doi: 10.1126/science.aao1887.

[26] Aaron P. Ragsdale, Timothy D. Weaver, Elizabeth G. Atkinson, Eileen G. Hoal, Marlo Möller, Brenna M. Henn, and Simon Gravel (May 2023). “A weakly structured stem for human origins in Africa”. Nature 617.7962, pp. 755–763. doi: 10.1038/s41586-023-06055-y.

[27] D. Reich, R. E. Green, M. Kircher, J. Krause, N. Patterson, E. Y. Durand, B. Viola, A. W. Briggs, U. Stenzel, P. L. F. Johnson, et al. (2010). “Genetic history of an archaic hominin group from Denisova Cave in Siberia”. Nature 468.7327, pp. 1053–1060.

[28] David Reich (2018). Who We Are and How We Got Here: Ancient DNA and the New Science of the Human Past. New York: Pantheon Books, 2018.

[29] Alan R. Rogers (2019). “Legofit: estimating population history from genetic data”. BMC Bioinformatics 20, p. 526. doi: 10.1186/s12859-019-3154-1.

[30] Alan R. Rogers (2022). “An efficient algorithm for estimating population history from genetic data”. Peer Community Journal 2, e32. doi: 10.24072/pcjournal.132.

[31] Alan R. Rogers, Nathan S. Harris, and Alan A. Achenbach (2020). “Neanderthal-Denisovan ancestors interbred with a distantly-related hominin”. Science Advances 6.8, eaay5483. doi: 10.1126/sciadv.aay5483.

[32] Arev P. Sümer, Hélène Rougier, Vanessa Villalba-Mouco, Yilei Huang, Leonardo N. M. Iasi, Elena Essel, Alba Bossoms Mesa, Anja Furtwaengler, Stéphane Peyrégne, Cesare de Filippo, Adam B. Rohrlach, Federica Pierini, Fabrizio Mafessoni, Helen Fewlass, Elena I. Zavala, Dorothea Mylopotamitaki, Raffaela A. Bianco, Anna Schmidt, Julia Zorn, Birgit Nickel, Anna Patova, Cosimo Posth, Geoff M. Smith, Karen Ruebens, Virginie Sinet-Mathiot, Alexander Stoessel, Holger Dietl, Jörg Orschiedt, Janet Kelso, Hugo Zeberg, Kirsten I. Bos, Frido Welker, Marcel Weiss, Shannon P. McPherron, Tim Schüler, Jean-Jacques Hublin, Petr Velemínský, Jaroslav Brůžek, Benjamin M. Peter, Matthias Meyer, Harald Meller, Harald Ringbauer, Mateja Hajdinjak, Kay Prüfer, and Johannes Krause (Feb. 2025). “Earliest modern human genomes constrain timing of Neanderthal admixture”. Nature 638.8051, pp. 711–717. doi: 10.1038/s41586-024-08420-x.

[33] P. J. Waddell (2013). “Happy New Year Homo erectus? More Evidence for Interbreeding with Archaics Predating the Modern Human/Neanderthal Split”. ArXiv 1312.7749. doi: 10.48550/arXiv.1312.7749.

[34] Peter J Waddell, Jorge Ramos, and Xi Tan (2011). “Homo denisova, correspondence spectral analysis, finite sites reticulate hierarchical coalescent models and the Ron Jeremy hypothesis”. ArXiv 1112.6424. doi: 10.48550/arXiv.1112.6424. eprint: 1112.6424.

[35] Duo Xu, Pavlos Pavlidis, Recep Ozgur Taskent, Nikolaos Alachiotis, Colin Flanagan, Michael DeGiorgio, Ran Blekhman, Stefan Ruhl, and Omer Gokcumen (2017). “Archaic hominin introgression in Africa contributes to functional salivary MUC7 genetic variation”. Molecular Biology and Evolution 34.10, pp. 2704–2715. doi: 10.1093/molbev/msx206.

[36] Yulin Zhang, Arjun Biddanda, Sarah Ariana Johnson, Colm O’Dushlaine, and Priya Moorjani (2026). “Recovering signatures of archaic introgression using ancestral recombination graphs”. bioRxiv. doi: 10.64898/2026.03.03.709416.

