## Supplementary materials for "The Genetic Legacy of Introgression from Two Deeply Divergent Lineages in the Genus *Homo*"

### Supplementary Materials for “Ancient Humans Interbred with Two Distinct Populations of Distant Relatives”

April 3, 2026

Fig. S1 shows the site pattern frequencies used in this analysis. They were generated by Legofit's `sitepat` program, which was run by the slurm script `sitepat.slr`. This script can be found in directory `data/xyrad` within the archived data and code. It also generates files of site pattern frequencies for bootstrap replicates.

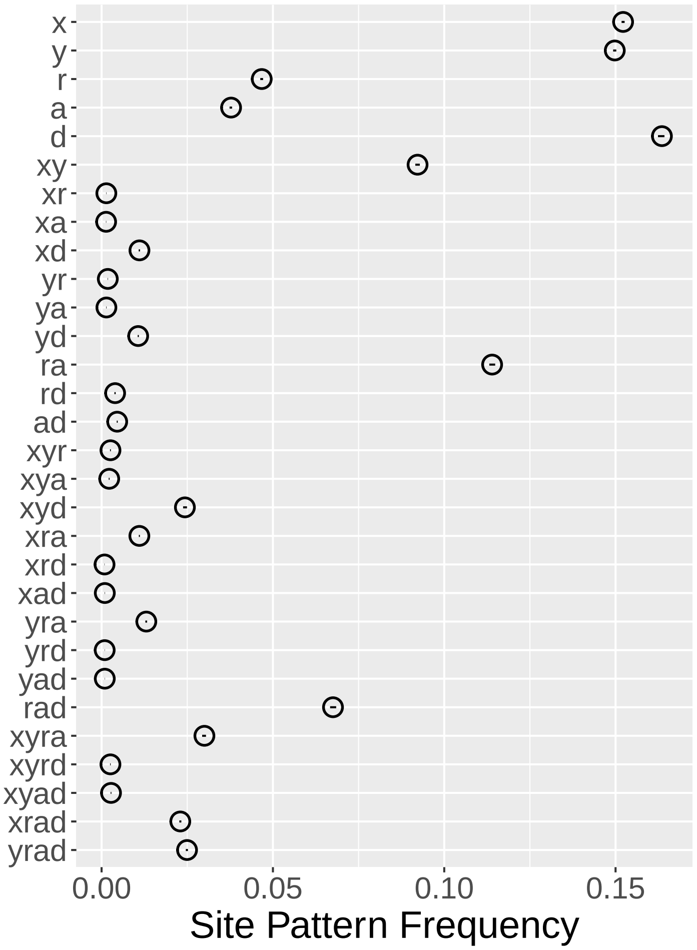


Figure S1: Site pattern frequencies. Horizontal lines (some of which are too small to see) are 95% confidence intervals made by moving blocks bootstrap.

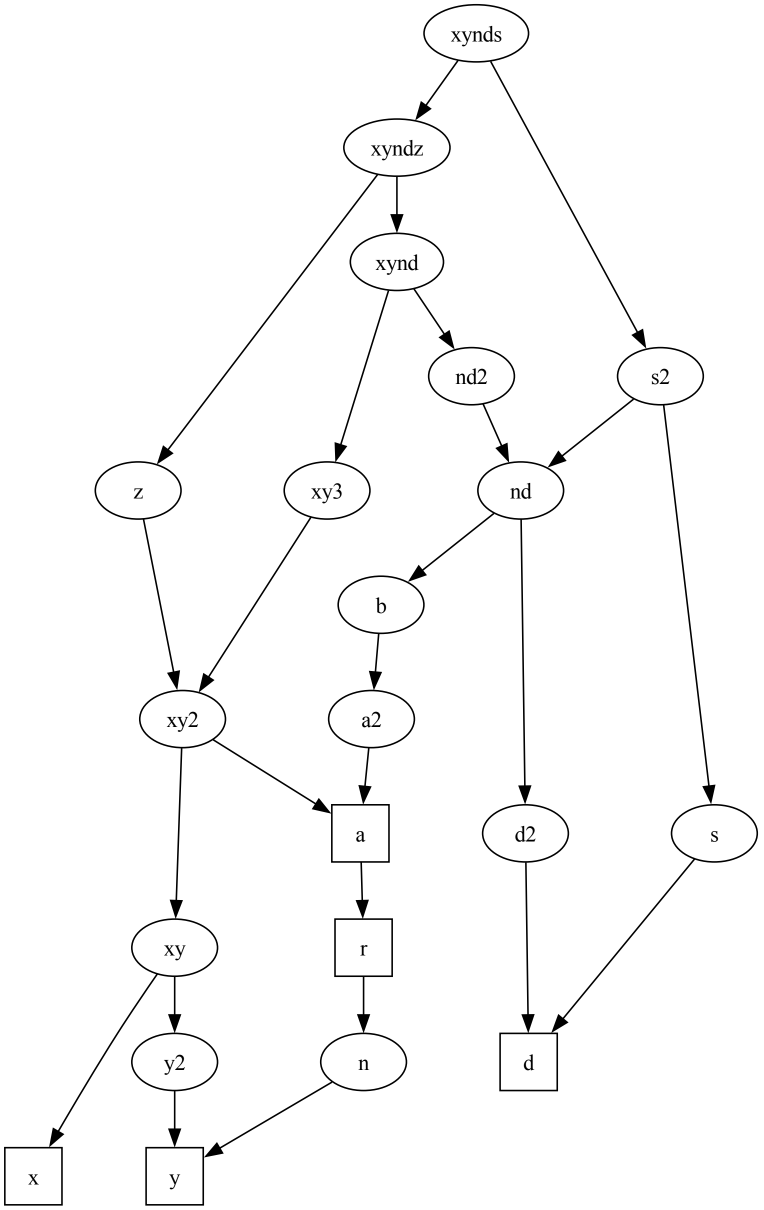


Figure S2: Legofit model $\alpha\beta\gamma\delta\zeta$

Fig. S2 shows a diagram of the network used in `Legofit` analysis of model $\alpha\beta\gamma\delta\zeta$. It was generated from file `a.lgo` within directory `data/xyrad/abcdz`. The latter file is the input used by `Legofit` to describe population history. To make a diagram of this model of history, the first step was to run `legosim -plot abcdz.dot a.lgo`, which generated `abcdz.dot`. Then, we used the `dot` program of Graphviz [1]: `dot -Tpdf abcdz.dot > abcdz.pdf`, which generated `abcdz.pdf`. We edited `abcdz.dot` by hand to adjust vertical spacing of some nodes.

Table S1: Model-averaged parameter estimates with 95% confidence intervals made by moving-blocks bootstrap [2]. Time unit is generations.

| | | 95% Confidence Bounds | |
|---|---|---|---|
| Parameter | Estimate | low | high |
| $m_\alpha$ | 0.018 | 0.016 | 0.019 |
| $m_\beta$ | 0.020 | 0.015 | 0.024 |
| $m_\gamma$ | 0.045 | 0.039 | 0.050 |
| $m_\delta$ | 0.058 | 0.049 | 0.070 |
| $m_\zeta$ | 0.196 | 0.124 | 0.264 |
| $T_A$ | 6210 | 6015 | 6387 |
| $T_B$ | 22507 | 21742 | 23164 |
| $T_D$ | 4331 | 4082 | 4547 |
| $T_{ND}$ | 25056 | 24856 | 25433 |
| $T_R$ | 3831 | 3633 | 3955 |
| $T_{XY}$ | 838 | 243 | 1505 |
| $T_{XYNDS}$ | 81836 | 76167 | 86407 |
| $T_{XYNDZ}$ | 44838 | 40738 | 49234 |
| $2N_B$ | 128629 | 97160 | 421263 |
| $2N_N$ | 7852 | 7623 | 7991 |
| $2N_{ND}$ | 1978 | 1100 | 2464 |
| $2N_S$ | 2431782 | 1903552 | 4974453 |
| $2N_{XY}$ | 48645 | 46521 | 51572 |
| $2N_{XYND}$ | 42171 | 41449 | 42795 |
| $2N_Z$ | 2853235 | 624208 | 5675017 |

Table S1 was generated in directory `data/xyrad`, using the command `bootci.py all.bma > all.bootci`. The input file, `all.bma` was generated by `booma */b2.bepe -F */b2.flat > all.bma`.

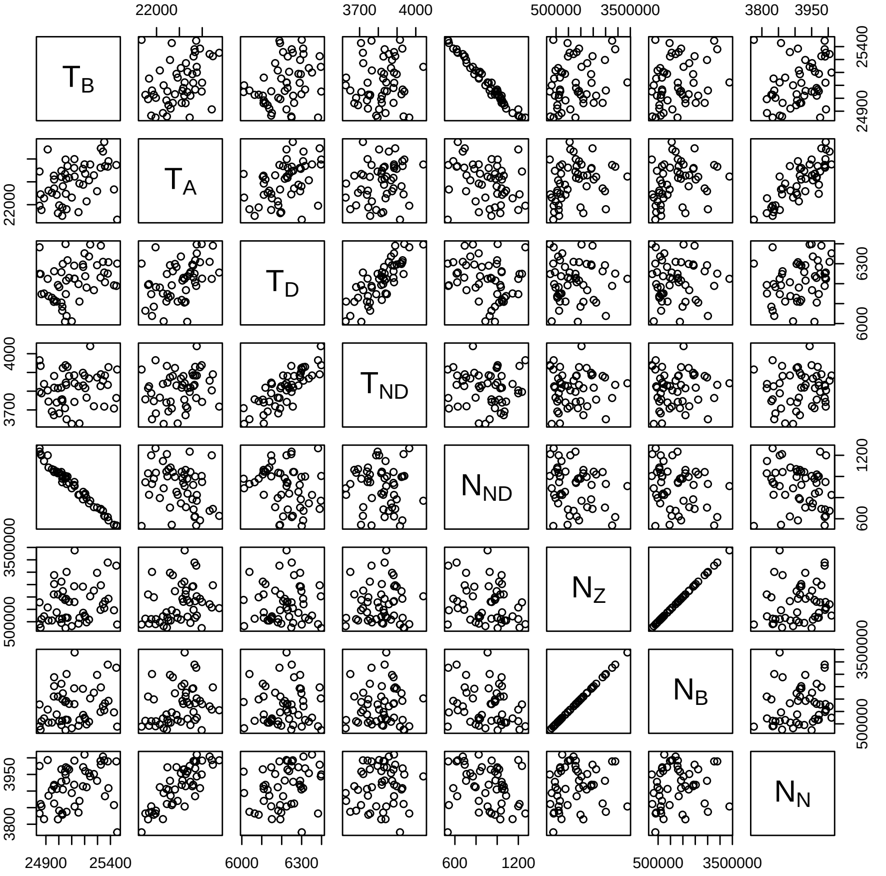


Figure S3: Scatterplot matrix showing associations between parameters. Each panel has 51 dots representing the real data and 50 bootstrap replicates. Plot includes only those parameters that correlate strongly (Pearson's $r \geq 0.8$) with some other parameter.

The scatterplot matrix in Fig. S3 shows associations between pairs of parameters. The tight correlations there imply identifiability problems. The figure was generated using the R script `b2pairs.r` within directory `data/xyrad/abcdz`.

Table S2: Estimates of $T_{XY}$ in years (assuming 29-y generations) under the model ($\alpha\beta\gamma\delta$) proposed by Rogers et al. [4]. Rows 1 and 2 are from Rogers et al.[4] and Rogers [3]. The last row is newly reported here.

| | 95% CI | | | Chagyrskaya |
|---|---|---|---|---|
| $\hat{T}_{XY}$ | lower | upper | algorithm | genome |
| 22,471 | 2,679 | 71,220 | stochastic | not used |
| 7,858 | 549 | 12,438 | deterministic | not used |
| 2,033 | 253 | 4,269 | deterministic | used |

In table S2, estimates of $T_{XY}$ get less and less plausible as we improve accuracy of calculations (row 2) and add data (row 3).

### References


[1] John Ellson et al. "Graphviz—open source graph drawing tools". In: *Graph Drawing*. Ed. by Petra Mutzel, Michael Jünger, and Sebastian Leipert. Berlin, Heidelberg: Springer, 2002, pp. 483–484. ISBN: 978-3-540-45848-7.

[2] Regina Y. Liu and Kesar Singh. "Moving blocks jacknife and bootstrap capture weak dependence". In: *Exploring the "Limits" of the Bootstrap*. Ed. by Raoul LePage and Lynne Billard. New York: Wiley, 1992, pp. 225–248.

[3] Alan R. Rogers. "An efficient algorithm for estimating population history from genetic data". *Peer Community Journal* 2 (2022), e32. DOI: 10.24072/pcjournal.132.

[4] Alan R. Rogers, Nathan S. Harris, and Alan A. Achenbach. "Neanderthal-Denisovan ancestors interbred with a distantly-related hominin". *Science Advances* 6.8 (2020), eaay5483. DOI: 10.1126/sciadv.aay5483.
